# A phylogenomic approach to resolving interrelationships of polyclad flatworms, with implications for life history evolution

**DOI:** 10.1101/2022.07.14.500079

**Authors:** Jessica A. Goodheart, Allen G. Collins, Michael P. Cummings, Bernhard Egger, Kate A. Rawlinson

**Author notes:** Co-senior authors (Bernhard Egger, Kate A. Rawlinson).

## Abstract

Platyhelminthes (flatworms) are a diverse invertebrate phylum that are useful for exploring life history evolution. Within Platyhelminthes, only two clades develop through a larval stage: free-living polyclads and parasitic neodermatans. Neodermatan larvae are considered evolutionarily derived, whereas polyclad larvae are hypothesized to be retained from the last common ancestor of Platyhelminthes – and Spiralia – due to ciliary band similarities among polyclad and other spiralian larvae. However, larval evolution has been challenging to investigate within polyclads due to low support for deeper phylogenetic relationships. To investigate polyclad life history evolution, we generated transcriptomic data for 21 species of polyclads to build a well-supported phylogeny for the group. We then used ancestral state reconstruction to investigate ancestral modes of development (direct vs indirect) within Polycladida, and flatworms in general. The resulting tree provides strong support for deeper nodes and we recover a new monophyletic clade of early branching cotyleans. Early branching clades of acotyleans and cotyleans possess diverse modes of development, suggesting a complex history of larval evolution in polyclads that likely includes multiple losses and/or multiple gains. Our ancestral state reconstructions in a previous platyhelminth phylogeny also suggests that similarities in larval morphology between flatworms and other phyla may have re-emerged secondarily or are convergently evolved.

## BACKGROUND

Flatworms (Platyhelminthes Minot, 1876) are among the most diverse invertebrate phyla, with an estimated 100,000 parasitic and free-living species [1]. They belong to the group of animals called Spiralia Schleip, 1929, which includes 12 other phyla; nemerteans, annelids, phoronids, ectoprocts, brachiopods, gastrotrichs, molluscs, entoprocts, chaetognaths, rotifers, micrognathozoans and gnathostomulids [2]. Many spiralians undergo indirect development whereby the embryo develops into the young adult form through a distinct larval stage. There are notable similarities in the structure and functional roles of larval characters amongst spiralians, such as ciliary bands for swimming and feeding [3], and these have led to hypotheses of their homology [4]. Two clades within the flatworms, Polycladida Lang, 1881 and Neodermata Ehlers, 1984, contain taxa with indirect development [5–8]. Although the complex life cycles of neodermatan flatworms and their larvae have been considered evolutionarily derived (i.e. an intercalated larval stage) [9], the biphasic life cycle of polyclads has been considered the ancestral condition for Platyhelminthes [9], and thought to be retained from the last common ancestor of the Spiralia due to similarities in larval ciliary bands and spiral cleavage patterns [4]. Recent phylogenomic analyses have, however, produced topologies for Platyhelminthes that call this hypothesis into question [10,11]. The position of the polyclads recovered in these analyses suggests that it is more parsimonious to view polyclad larvae as one or more independent acquisitions limited to this group [11].

Polyclads (order Polycladida) offer an interesting group within which to explore the evolution of life history strategies and larval characters. They are a clade of marine flatworms that, as adults, are generally found on the seafloor in coastal habitats (Figure 1A), but they have also been collected in the deep sea and in the water column [12,13]. The most common mode of development in Polycladida might be direct [14], whereby the hatchling resembles a sexually immature adult worm, i.e. dorso-ventrally flattened (Figure 1B-D), with a uniform covering of motile cilia. However, there are many species that develop indirectly, through a planktonic phase that has transient larval features which make it morphologically distinct from the adult form. These larval features include lobes, or protrusions, upon which are bands of longer motile cilia used for swimming in the water column (Figure 1 E-K). These features are lost at metamorphosis and the transition to a benthic niche [8,15]. The Polycladida has traditionally been divided into two suborders based on multiple morphological characters of the adult worms, including the presence (Cotylea Lang, 1884, ~350 species) or absence (Acotylea Lang, 1884, ~450 species) of a ventral adhesive structure [16–18], known as a cotyl [19,20]. Most cotyleans examined to date hatch as an 8-lobed larval stage (known as Müller’s larva) (e.g. Figure 1 E,F & H). There are four known exceptions; *Prosthiostomum acroporae* (Rawlinson et al., 2011) undergoes intra-capsular metamorphosis – reabsorbing its 8 larval lobes before hatching (intermediate development) [21] (Figure 1G), *Pericelis cata* Marcus & Marcus, 1968 exhibits poecilogony (i.e., hatchlings emerge from an egg plate with and without larval characters) [22], *Boninia divae* Marcus & Marcus, 1968 hatchlings have reduced lobes [22], and *Theama mediterranea* Curini-Galletti et al., 2008 hatches as juveniles without lobes and ciliary bands [23]. By contrast, taxa assigned to Acotylea mainly exhibit direct development, but species with intermediate and indirect development do exist, with a diversity of larval morphologies (e.g., 6- and 8-lobed Müller’s larvae, 4-lobed Götte’s larva (Figure 1I) and an 8-lobed, dorso-ventrally flattened Kato’s larva [14]). Larvae with 10 lobes have also been described [9,14,24](Figure 1J,K).

**Figure 1.**
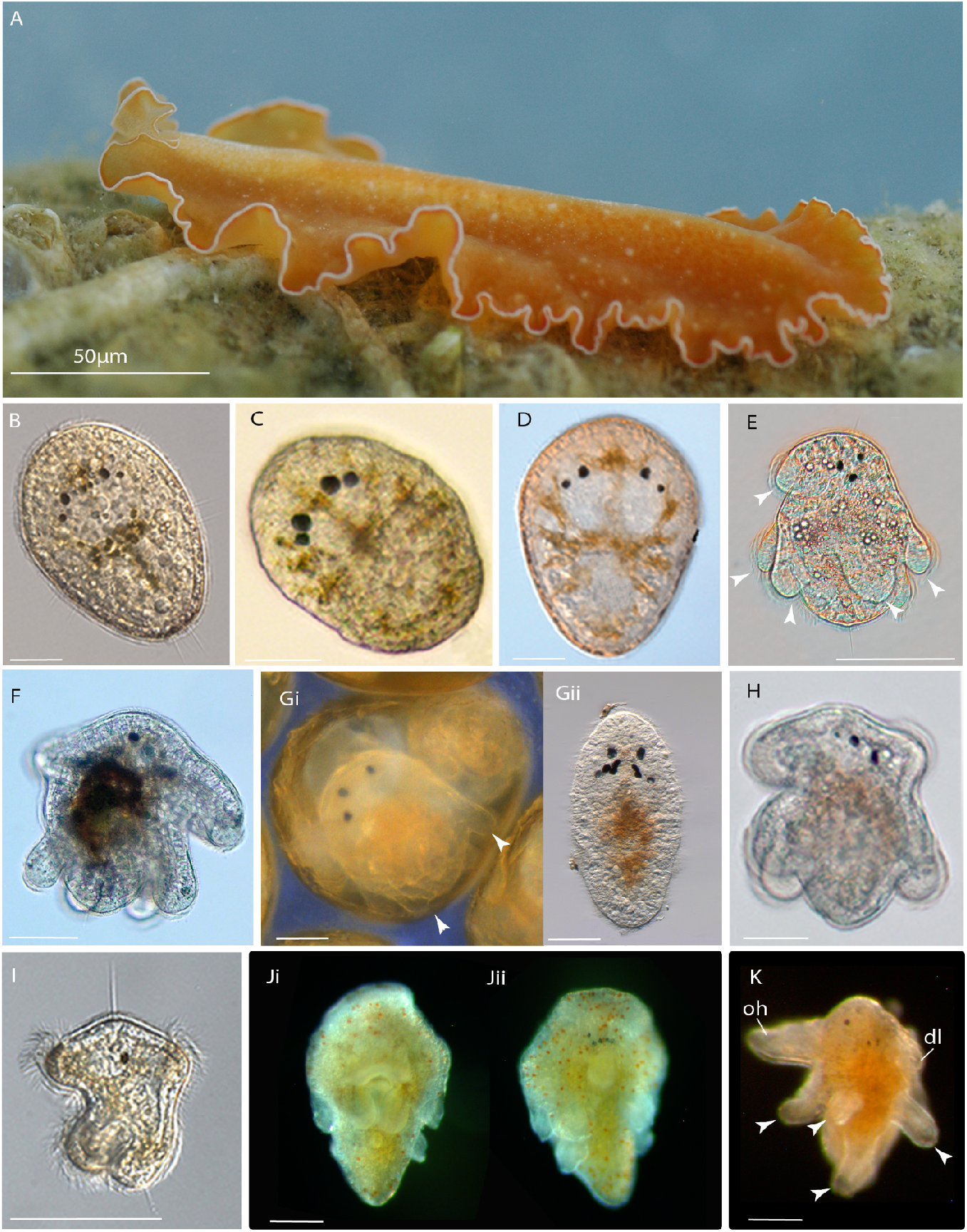
Polyclad flatworms are dorso-ventrally flattened as adults, some species develop directly into this body plan, others develop indirectly through larval forms with transient features such as lobes and ciliary tufts and bands. A) Adult polyclad, *Yungia* sp. B-D) Hatchlings of direct developing species; *Euplana gracilis* (B), *Notocomplana* sp. (Schmarda, 1859) (C) and *Echinoplana celerrima* Haswell, 1907(D). E-K) Hatchlings of indirect developing taxa; newly hatched Müller’s larva of *Cycloporus gabriellae* Marcus, 1950, showing long motile cilia on lobes (arrowheads) (E), newly hatched Müller’s larva of *Prosthecereaus crozieri* (Hyman, 1939) (F); *Prosthiostomum acroporae* shows ‘intermediate development’ with the embryo developing 8 larval lobes (arrowheads) inside the egg capsule (G i), but most individuals undergo intra-capsular metamorphosis – reabsorbing larval lobes before hatching (G ii); Müller’s larva of *Prosthiostomum siphunculus* (Delle Chiaje, 1822) (H), four-lobed Götte’s larva of *Stylochus ellipticus* (Girard, 1850) (I), ventral (Ji) and dorsal (Jii) view of 10-lobed larva collected from the plankton, species unknown; and lateral view of 10-lobed larva collected from the plankton (K). B-K scale 50 μm. Species in B-H are sequenced in this study.

Recent molecular phylogenies of the Polycladida have significantly increased our understanding of the interrelationships within the order [20,25–34]. However, these studies used one or a few genes for their inferences, primarily 18S, 28S and COI. It is well known that phylogenetic inferences using single gene data matrices may represent limited approximations of the relationships among taxa (as they represent gene trees as opposed to species trees) [35] and are subject to limitations in scope due to differences in rates of evolution across genes (e.g., [36]). They also offer a comparatively low number of phylogenetically informative positions. In the case of polyclads, these challenges have led to phylogenies that are often well-supported near the tips but provide lower resolution of earlier branching lineages in the polyclad tree.

The two goals of this paper were to generate a well-supported phylogenetic hypothesis for Polycladida inferred from transcriptomic data; and to use this framework to assess the evolutionary origin of a larval stage (i.e., indirect development) in polyclads. The use of transcriptomes for phylogeny removes the potential bias related to selecting specific genes for analysis *a priori*. We generated RNAseq data from 21 polyclad species and developed a data matrix that included a further six transcriptomes for polyclads, and 9 transcriptomes for non-polyclad flatworms. We scored the mode of development for these taxa from the literature and personal observation. Here, indirect development was scored if the species has distinct and transient characters that are not retained in the adult body plan (i.e., larval characters, specifically lobes and/or ciliary bands/plates/tufts), whereas direct development was called if no larval characters are recorded during embryogenesis and at hatching. We used the phylogenomic relationships to reconstruct the ancestral mode of development among the polyclad lineages in our tree to discern where particular development modes originated. Finally, we used a previous phylogenomic-based tree of the phylum to infer the ancestral mode of development at nodes within the flatworm phylogeny. This enabled us to determine whether it is phylogenetically congruent to consider indirect development (and characters, such as larval ciliary bands) homologous among polyclad families and suborders, and among different flatworm orders (i.e., polyclads and neodermatans). This work represents an important step in our comprehension of polyclad phylogeny and life history evolution among flatworm clades.

## RESULTS

### Assembly and data matrix properties

Our final data matrix includes transcriptomes of 27 polyclad species from at least 23 genera (Table S1), plus nine outgroup taxa representing other platyhelminth lineages (Catenulida Meixner, 1924, Macrostomorpha Doe, 1986, Gnosonesimida Karling, 1974, Cestoda Gegenbaur, 1859, and Tricladida Lang, 1881) [11]. Once assembled, the number of contiguously assembled sequences (“contigs”) per sample ranged from 62,613 (*Xenoprorhynchus* sp. I Laumer & Giribet, 2014) to 819,086 (*Prostheceraeus vittatus* (Montagu, 1815)) (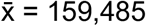; Table S2). N50 ranged from 520 bp (*Xenoprorhynchus* sp. I) to 2,041 bp (*Boninia divae*) (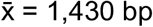; Table S2), and Benchmarking Universal Single-Copy Orthologs (BUSCO) scores (compared to the metazoa_odb10 database) ranged from 28.8% complete (Xenoprorhynchus sp. I) to 91.4% complete (Prosthiostomum siphunculus) with all new polyclad transcriptomes surpassing 83% complete. Our data matrix consisted of 4,469 orthologous groups and 5,081,724 nucleotide positions (58.0% complete with 0.32% ambiguous characters; Table S2). Individual species ranged from 848,391 bp (16.7% of full alignment length, *Xenoprorhynchus* sp. I) to 3,916,323 bp (77.1% of full alignment length, *Leptoplana tremellaris* (Müller OF, 1773)), with an average of 2,947,307 bp across all species (Table S2).

### Phylogenetic analyses

The inferred topologies were identical across all three analyses (with some variation in branch lengths), with 100% bootstrap support for all branches in our analysis accounting for heterotachy. Here we report results only for our maximum likelihood phylogeny of Polycladida constructed using RAxML-NG with our matrix partitioned by codon position (Figure 2). Results from the other two analyses are provided in our dryad repository (https://doi.org/10.6076/D1JG60). All 20 search replicates in our partitioned analysis returned the same best tree topology (Figure 2) and[37] indicate strong support for the monophyly of Polycladida (bootstrap support (BS) = 100) and the taxonomic sub-clades Cotylea *sensu* Bahia et al. 2017 (BS = 100) and Acotylea (BS = 100) *sensu* Dittmann et al. 2019. The enigmatic genera *Cestoplana* Lang, 1884, *Pericelis* Laidlaw, 1902, *Boninia* Bock, 1923, and *Theama* Marcus, 1949 fall within the Cotylea, and form a novel clade, clade 1 (BS = 100; Figure 2), that is sister to the rest of cotyleans (BS = 100). Within clade 1, Boniniidae Bock, 1923 and Theamatidae Marcus, 1949 are sister taxa, supporting the recently proposed superfamily Boninioidea Bock, 1923 [28]. The cotylean families Prosthiostomidae Lang, 1884, Pseudocerotidae Lang, 1884, and Euryleptidae Stimpson, 1857, are recovered with high support (all with BS = 100).

**Figure 2.**
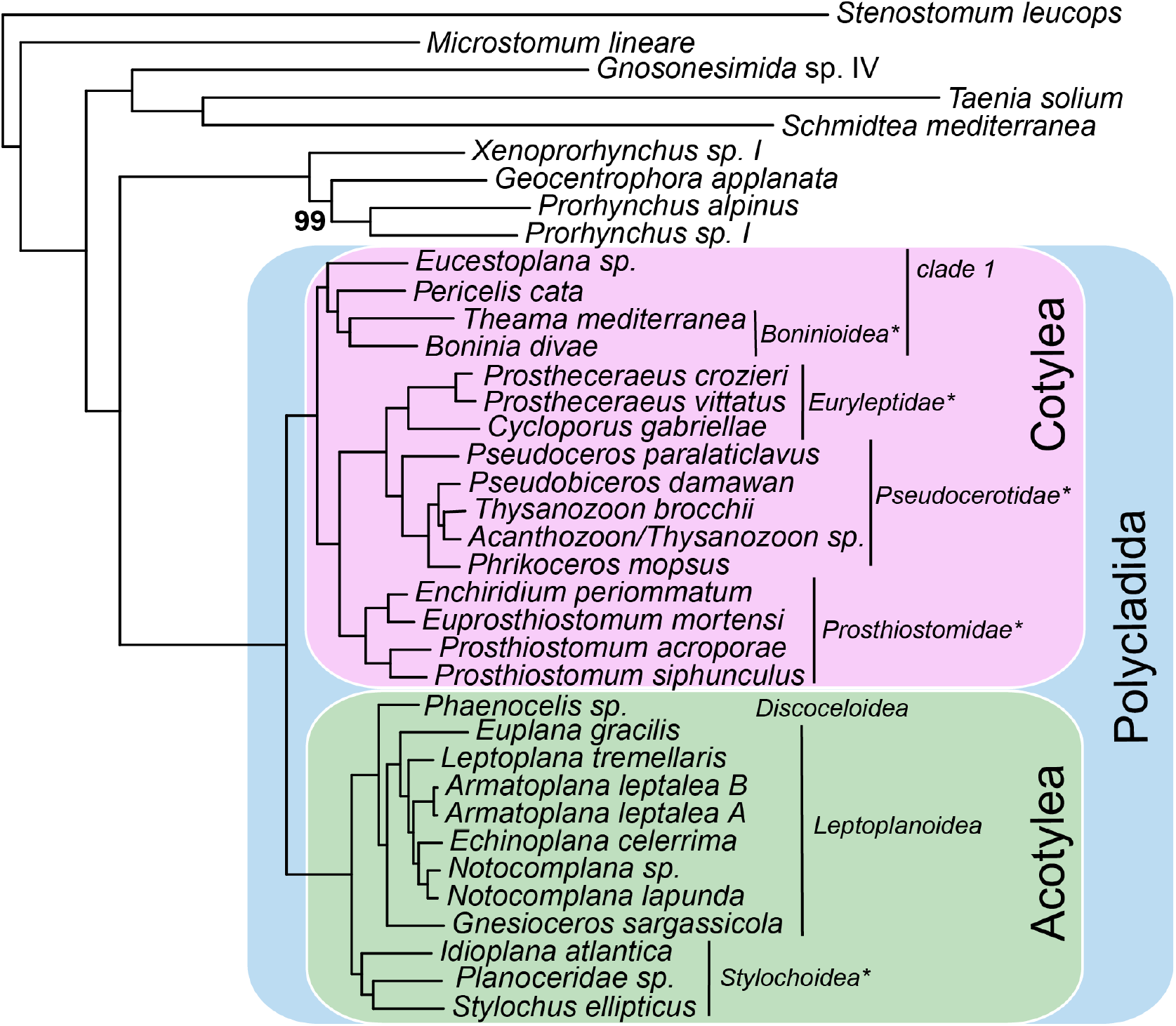
Maximum likelihood phylogeny of Polycladida constructed using RAxML-NG from a concatenated nucleotide matrix of 4,469 genes partitioned by codon position. Unless indicated otherwise, all branches have 100% bootstrap support. Each branch also has 100% bootstrap support in our IQ-TREE analysis accounting for heterotachy. The blue box indicates the clade Polycladida, and the pink and green boxes indicate the clades for Cotylea and Acotylea, respectively. Superfamily and family-level systematic affinities supported by this analysis are marked with an asterisk.

In Acotylea, there are three superfamilies: Stylochoidea Stimpson, 1857, Leptoplanoidea Ehrenberg, 1831 and Discoceloidea Laidlaw, 1903. Our results are consistent with a monophyletic Stylochoidea *sensu* Dittmann et al., 2019 (BS = 100), and its position as the sister lineage to the rest of the suborder [20,27,28]. Leptoplanoidea *sensu* Dittmann et al., 2019 is rendered paraphyletic by the inclusion of *Euplana gracilis* Girard, 1853 (BS = 100) (Table S1). As a single undescribed species of *Phaenocelis* Stummer-Traunfels, 1933 is the only confirmed taxon of Discoceloidea included in our analysis, support for this superfamily could not be assessed.

The family Gnesiocerotidae Marcus & Marcus, 1966 (*sensu* Faubel 1983) represented here by *Gnesioceros* Diesing, 1862 and *Echinoplana* Haswell, 1907, was not supported (Table S1). The newly created family Notocomplanidae Litvaitis et al., 2019 was monophyletic. The families Stylochidae Stimpson, 1857, Idioplanidae Dittmann et al. 2019, Cryptocelidae Laidlaw, 1903, Euplanidae Marcus & Marcus 1966, are represented by only one species in our analysis, therefore monophyly of these families could not be assessed.

### Ancestral character estimation

Mode of development is unknown for 11 of the 36 species (31%) in our phylogeny (Table S3). However, for 6 of these 11 species, there is developmental information for one or more congeneric species. Therefore, we coded the mode of development at the genus-level for all taxa in our inferred polyclad phylogeny (except an undescribed Planoceridae species that we coded at the family level). Our ancestral state reconstruction shows indirect development at the base of a clade of cotyleans that includes the families Prosthiostomidae, Euryleptidae and Pseudocerotidae; and direct development at the base of a clade of acotyleans that includes members of the superfamilies Discoceloidea and Leptoplanoidea (Figure 3). However, our ancestral state reconstruction was unable to confidently reconstruct development type at deeper nodes in the polyclad tree; i.e., the base of cotylean clade 1 (scaled likelihoods: indirect = 0.865, direct = 0.135), Cotylea (scaled likelihoods: indirect = 0.874, direct = 0.126), Stylochoidea (scaled likelihoods: indirect = 0.547, direct = 0.453), Acotylea (scaled likelihoods: indirect = 0.488, direct = 0.512), and Polycladida (scaled likelihoods: indirect = 0.659, direct = 0.341). The early branching cotylean clade 1 and acotylean Stylochoidea show the greatest diversity in mode of development among the polyclads. Due to sampling bias in favor of polyclad species, we were also unable to reconstruct the ancestral mode of development at deeper nodes within Platyhelminthes. To account for this bias, we also reconstructed ancestral mode of development using a previously published phylogeny of flatworms [[11]. In this analysis, the ancestor of Polycladida and Prorhynchida Karling, 1974, was reconstructed as a direct developer (scaled likelihoods: indirect = 0.093, direct = 0.908), as were most other ancestral nodes within Platyhelminthes (Figure 4). Together, these two ancestral state reconstructions indicate that indirect development in polyclads may have evolved in the polyclad stem lineage and been lost in the Discoceloidea and Leptoplanoidea and *Theama* in Cotylea clade 1. However, our results could also indicate multiple gains of indirect development in polyclads (i.e. at the ancestral node of the Euryleptidae/ Pseudocerotidae/ Prosthiostomidae clade and in Planoceridae).

**Figure 3.**
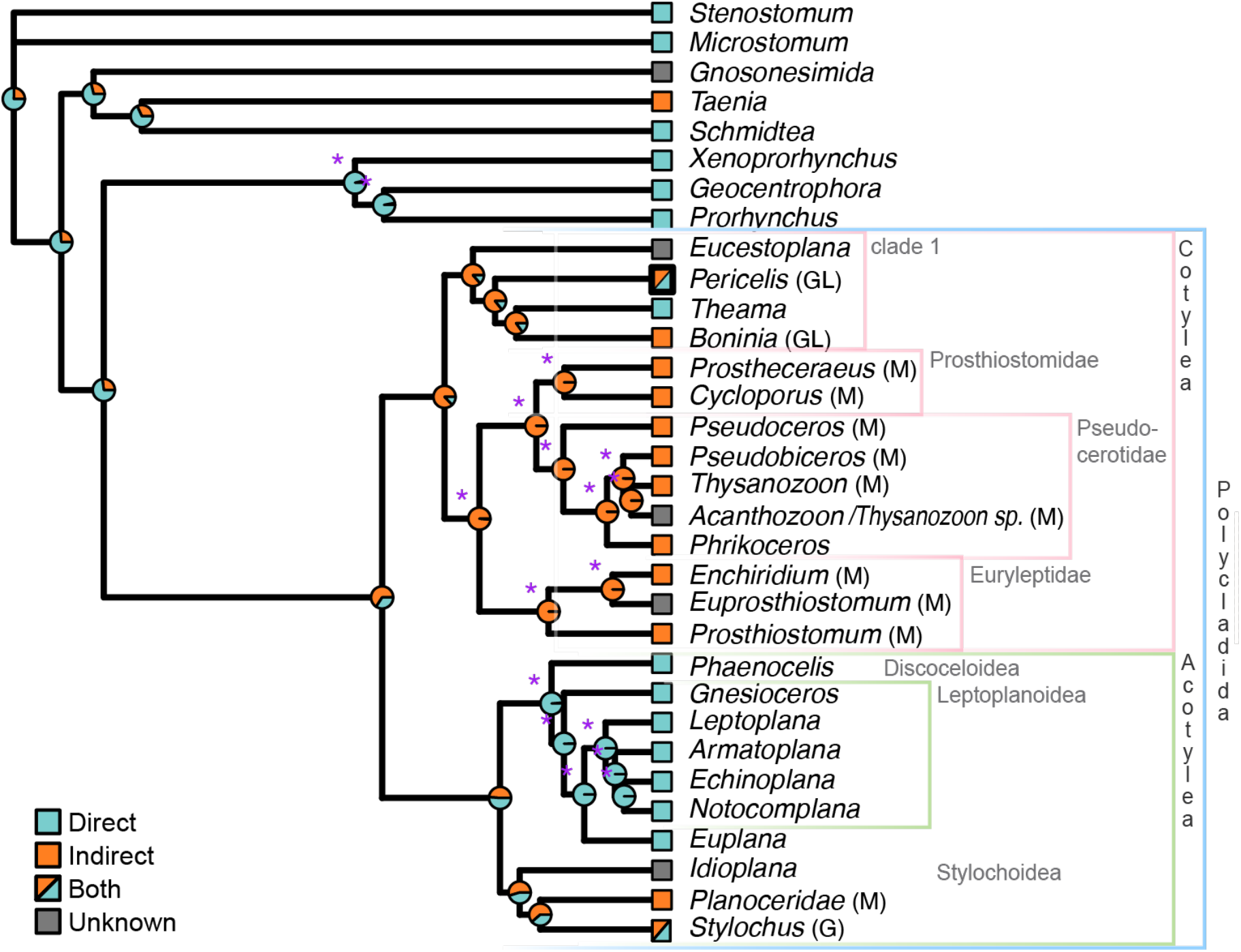
Ancestral state reconstruction analysis in Polycladida for the presence of indirect (orange), direct (blue) development, or both strategies in the genus (orange/blue; thin border denotes species within the genus have both strategies; thick border denotes some species within the genus exhibits poecilogony). Pie charts on the nodes are scaled marginal likelihoods calculated using the ace function in APE. The purple asterisks denote significant nodes as defined by proportional likelihood significance tests [38] with a likelihood difference of at least 2 or higher. Type of larva in brackets after genus name; M = Müller’s, G = Götte’s and GL = Götte’s-like.

**Figure 4.**
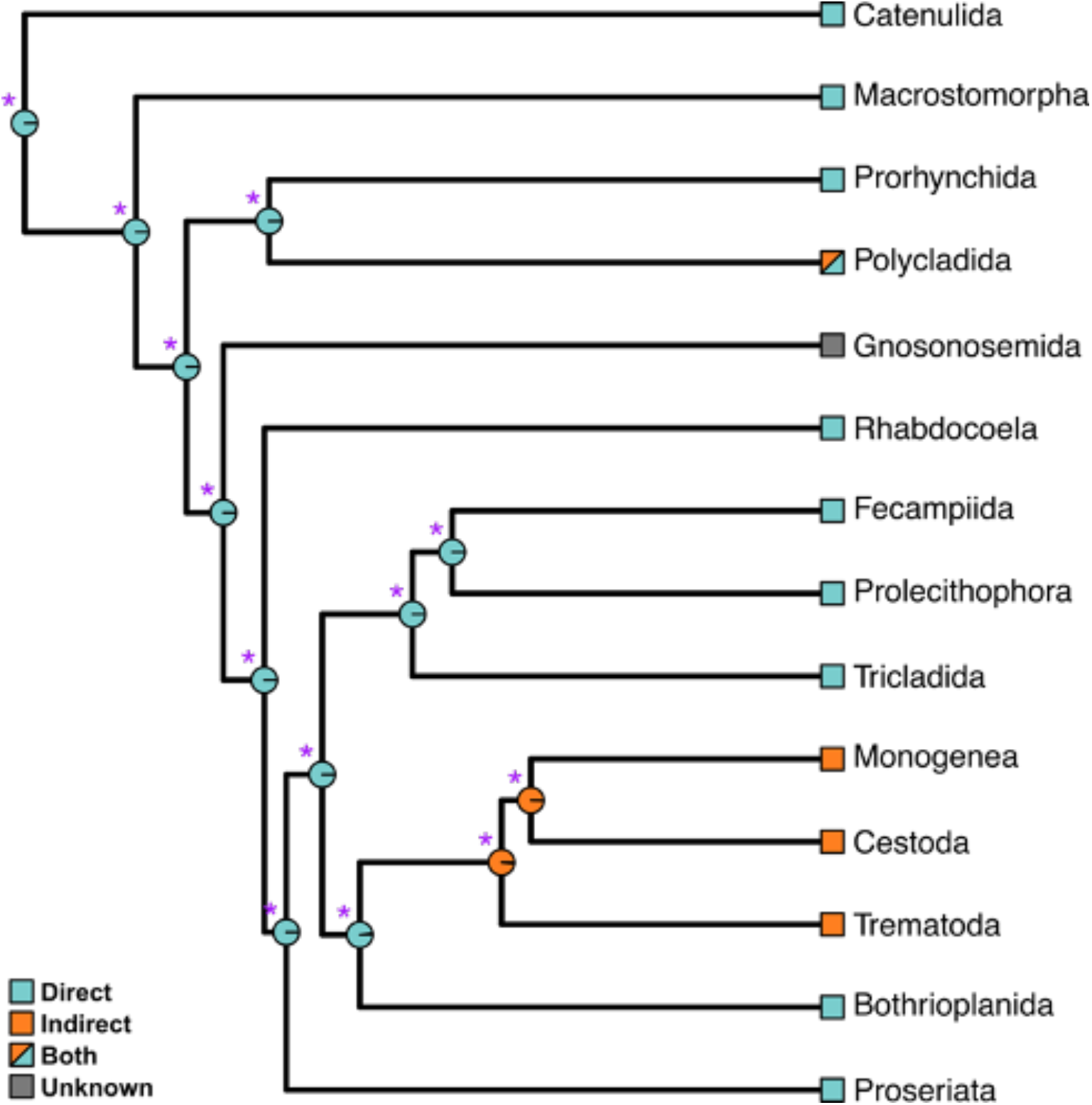
Ancestral state reconstruction analysis in the phylum Platyhelminthes for the presence of indirect (orange), direct (blue) development or both strategies in the order (orange/blue). Pie charts on the nodes are scaled marginal likelihoods calculated using the ace function in APE. The purple asterisks denote significant nodes as defined by proportional likelihood significance tests [38] with a likelihood difference of at least 2 or higher.

## DISCUSSION

In this study we have significantly increased the breadth of RNAseq sampling in Polycladida to generate a robust phylogenetic hypothesis for this group and provide a framework to investigate the evolution of modes of development in polyclads. We have constructed a massive data matrix (> 5 million nucleotide positions; 4,469 genes) for inferring the phylogeny of Polycladida, which has provided very high confidence in the phylogenetic inferences presented here (bootstrap support (BS) = 100 within Polycladida and BS > 98 across outgroup branches). This success implies that transcriptomic data will be particularly useful for resolving the phylogeny of Polycladida moving forward.

### The phylogeny of Polycladida

Analysis of this large dataset has resulted in a tree that shows some correspondence with single- and few gene-based trees, and importantly has resolved uncertainties around early branching lineages in Polycladida (Figure 2). Our phylogeny shows full support at all nodes within Polycladida, including deeper nodes that in past studies have often had weak or no support [20,27,28], though not exclusively [33]. We have not modified the nomenclature of clades in Polycladida based on our phylogeny due to the limited taxonomic coverage, but we discuss the implications of the tree topology for systematics and highlight taxa to include in future transcriptome-based analyses. We also discuss limited changes to the classification of some species based on our phylogeny. If not otherwise indicated, our nomenclature and taxonomy definitions follow Dittmann et al. [27].

#### Cotylea

One novel finding resulting from our analysis is a new monophyletic clade (clade 1, Figure 2) sister to all other Cotylea that includes the families Cestoplanidae Lang, 1884, Theamatidae, Boniniidae and Diposthidae Woodworth 1898 *sensu* Litvaitis et al. 2019. Morphology based classifications placed Cestoplanidae and Theamatidae within Acotylea [17,18] due to the lack of a ventral cotyl [20]. Recent molecular phylogenies supported the Cestoplanidae Lang, 1884 and Theamatidae as early branching members of Cotylea [20,27,28,33]; with Cestoplanidae either as the sister group of all remaining cotyleans [20,27,28,33], or as sister group of Diposthidae, together forming the sister group of all remaining cotyleans [20,38]. Our analysis places Cestoplanidae within a new clade, with Boniniidae and Diposthidae. In future analyses, the inclusion of other cotyleans identified as early branching (e.g., species belonging to *Anonymus* Lang, 1884, *Chromoplana* Bock, 1922, and *Chromyella* Correa, 1958 [20,27,28]) will be important to see if these species form part of clade 1 and whether clade 1 remains the sister to the rest of Cotylea.

In our reconstruction, *Pseudoceros* Lang, 1884 is the sister group of all other pseudocerotids, reflecting topologies of single- and few-gene trees [26–28,33]. The family Pseudocerotidae contains genera with single or duplicated male copulatory organs. Genera with duplicated male organs (e.g., *Thysanozoon* Grube, 1840, *Pseudobiceros* Faubel, 1984) were supported as a monophyletic cluster in the phylogenetic reconstruction of Litvaitis et al. [20], although genera with single and duplicated genital organs were intermingled in other studies [20,26–28,33]. Here, *Thysanozoon brocchi* (duplicated male apparatus) is sister group of ‘*Acanthozoon* or *Thysanozoon* sp.*’*, where *Acanthozoon* Collingwood, 1876 features a single male apparatus. Although our species determination did not allow distinguishing between *Acanthozoon* and *Thysanozoon*, the hypothesis of a single origin of duplicated male organs put forward by Litvaitis et al. [20] suggests the sample labeled ‘*Acanthozoon*/*Thysanozoon* sp.’ in Figs. 2–3 is a *Thysanozoon* rather than an *Acanthozoon*.

Further taxon sampling is required to resolve the relationships within the families Prosthiostomidae and Euryleptidae. Within the Prosthiostomidae, *Euprosthiostomum* Bock, 1925 is sister group of *Enchiridium* Bock, 1913 (this study), although in two recent studies, *Euprosthiostomum* was recovered as sister group of *Prosthiostomum* Quatrefage, 1845 [20,34]. All three studies used the same species of *Euprosthiostomum*, *E. mortenseni* Marcus, 1948, and recovered the conflicting topologies with high support. Although our tree recovers a monophyletic Euryleptidae consistent with Tsunashima et al. [26], other recent phylogenies recovered paraphyletic Euryleptidae [20,27,28,33], which may be attributable to increased taxon coverage, i.e., inclusion of members of the family Stylostomidae Dittmann et al., 2019.

#### Acotylea

The interrelationships of the three acotylean superfamilies recovered here – Stylochoidea as sister to Discoceloidea + Leptoplanoidea, has been proposed by earlier studies [20,26–28,32,33,38]. However, our analysis renders the Leptoplanoidea *sensu* Faubel, [17,18] paraphyletic by the inclusion of *Euplana gracilis* within this clade. *E. gracilis* is the first member of Euplanidae to be included in a molecular phylogeny of polyclads. In Faubel’s system, the Euplanidae belong to the now suppressed superfamily Ilyplanoidea Faubel, 1983, [17,18], and later redefined as Discoceloidea [28]. Here, we transfer the family Euplanidae to the superfamily Leptoplanoidea.

Within Leptoplanoidea our analysis supports the paraphyly of the family Gnesioceridae Marcus & Marcus, 1966 found in recent studies [30–32,38]. *Gnesioceros sargassicola* (Mertens,1833) is recovered as the sister taxon to all other Leptoplanoidea [20,31,38], but a second species previously assigned to Gnesioceridae, *Echinoplana celerrima* Haswell, 1907, forms a derived clade with *Notocomplana* species. The sister relationship between *Echinoplana* and *Notocomplana* has also been recovered in Bahia et al. *[27]*. Unlike Notocomplanidae Litvaitis et al. 2019, which are diagnosed by the absence of a sclerotised stylet on the penis [20], the penis of *Echinoplana* is armed with numerous spines and hooks. Together these data suggest that *Echinoplana* does not belong to Gnesiocerotidae, but may form a new family closely related to Notocomplanidae, or be included in Notocomplanidae (thus prompting a renaming of the family to Echinoplanidae, as the name *Echinoplana* precedes *Notocomplana*). Including other gnesiocerotids (e.g., *Styloplanocera* Bock, 1913 and *Planctoplanella* Hyman, 1940) in future phylogenomic analyses will determine the valid members of this family.

Our analysis provides a phylogenetic foundation upon which future phylo-transcriptomic and genomic studies may build. The majority of families and genera, erected on a morphological basis by Faubel [17,18] and Prudhoe [16,18], have not been tested here (nor in any one molecular framework) leaving both the monophyly of families, as well as their inclusion in superfamilies, unresolved. We included species from 10 of 28 acotylean families and 5 of 15 cotylean families after Faubel [17,18], and 7 of 18 acotylean families and 5 of 10 cotylean families after Prudhoe [16]. The challenge therefore remains to increase taxonomic sampling in order to establish a new phylogenomic system which resolves the conflicting topologies and classifications based on morphology [16–18,39] and on single- or a few-genes [20,26–28].

### Life history evolution within Polycladida and Platyhelminthes

Our ancestral state reconstruction highlights the evolutionary complexity of mode of development within the Polycladida (Fig. 3). Our inability to significantly reconstruct the deeper nodes <u>in polyclads</u> is partly due to missing data, but also because early branching clades in both suborders contain taxa with different modes of development.

Although the vast majority of Acotylea studied to date have direct development, larvae are known to occur in two of the three current superfamilies; in Stylochoidea (in the closely related genera *Hoploplana*, *Planocera*, *Stylochus* and *Imogine*, summarized in [14]) and, less frequently, in Leptoplanoidea (*Notoplana australis* (Schmarda, 1859) [39] and possibly *Stylochoplana maculata* Quatrefage, 1845 [40]). An especially interesting case of larval evolution is found in the genus *Planocera*, where *Planocera reticulata* features a unique eight-lobed, multi-eyed and dorsoventrally flattened Kato’s larva but the congener *Planocera multitentaculata* develops via an eight-lobed, three-eyed, spherical Müller’s larva very similar to prosthiostomid, euryleptid and pseudocerotid Müller’s larvae [41,42]. Litvaitis *et al*. [20] suggested the presence of larvae could be a synapomorphy for Stylochoidea, but went on to add that because Stylochoidea (*sensu* Poche[43]) has been identified as the most early diverging lineage in Acotylea [20,27,28], it is more likely that larvae are a symplesiomorphy retained from the polyclad ancestor. Our phylogenetic analysis is consistent with Stylochoidea as the sister group of all other acotyleans, and our ancestral state reconstruction does not rule out the possibility that indirect development in the Stylochoidea is conserved from an ancestral polyclad. However, descriptions of development are missing in at least one stylochoidean genus in our analysis (*Idioplana*), which may impact our inferences. Furthermore, adding transcriptome and development data from a Latocestidae species, an unrepresented family of the Stylochoidea in this study, may help resolve this node.

Despite most cotylean taxa having indirect development, there was no significant support for this mode of development in the ancestral cotylean. This is likely due to the diversity of development recorded in the early branching clade 1, which may also impact inferences in Acotylea. Within this clade nothing is known about development of the cestoplanids, and the other members exhibit poecilogony (*Pericelis cata*, [22]), indirect (*Boninia divae*, [22]) and direct development (*Theama mediterranea*, [23]). Interestingly though, there are a number of similarities in the development of these three species; the hatchlings have just one eyespot, compared to two or more in other polyclad hatchlings, the egg capsules contain multiple embryos [22,23], and the larvae of *Pericelis cata* and *Boninia divae* have been described as atypical, with a smaller number of lobes that are reduced in size, more similar to the Götte’s larvae of acotylean species than the Müller’s larvae of other cotyleans [22]. It is possible that these modes of development represent intermediate evolutionary stages between acotylean direct development and the derived cotylean indirect development, or indirect development in both clades.

There are three possible evolutionary scenarios for indirect development in polyclads based on our findings; the first is that a Müller’s type larva could be the ancestral polyclad condition, which has been reduced to Götte-like larvae in the stylochids and cotylean clade 1 and lost completely in most Leptoplanoidea + Discoceloidea. The second, is that indirect development via a Götte-like larvae (as found in some acotyleans and also in some cotylean clade 1 taxa) may be the ancestral condition in polyclads, from which Müller’s larvae have independently evolved several times (in *P. reticulata* and in most cotyleans). Thirdly, instead of an ancestral polyclad larva, Müller’s and Götte’s larvae could be evolutionarily derived larval types within the polyclads that have each evolved several times independently. Although our present analyses cannot shed light on which scenario is more likely, they provide the framework for future analyses and highlight key taxa for further investigations into the mode of development, i.e. *Idioplana atlantica* (Bock, 1913) and Cestoplanidae.

Our second analysis reconstructed the ancestral development type for the whole phylum (Figure 4) using a previously published phylogeny [11]. This second analysis allowed us to infer the ancestral development mode for Polycladida and its sister lineage, Prorhynchida, because we reduced the polyclad-heavy sampling bias and limited outgroup sampling that led to lack of support for reconstructions in our polyclad phylogeny (Figure 3). This analysis points to polyclad larval stages being intercalated into the life cycle of a direct developing prorhynchid/polyclad ancestor along the polyclad stem lineage or within polyclads. Recent phylogenomic studies of flatworms have suggested it is more parsimonious to view polyclad larvae as one or more independent acquisitions limited to this group [10,11], and our analysis supports this.

These interpretations bring into question the hypothesis that indirect development in polyclads is retained from a spiralian ancestor with a biphasic life cycle and trochophore-like larva. This is a long-standing idea based on morphological similarities in larval stages (e.g. larval lobes and ciliary bands) and supported by the presence of spiral cleavage in polyclads and other non-flatworm spiralians [9,41,42], and similar embryonic origins of polyclad and spiralian trochoblasts [44]. Our analyses, instead, suggest that similarities in larval characters of polyclad and trochophore larvae may not be phylogenetically congruent, and may have evolved convergently. Another interpretation is that some elements of indirect development in a spiralian ancestor may have been secondarily derived in polyclads using some combination of novel and conserved developmental pathways. However, to reconstruct the developmental mode of the ancestral flatworm, and inform on homology of polyclad and trochophore larvae, future phylogenetic and ancestral state reconstruction analyses would require the inclusion of many more non-polyclad flatworms and multiple species from sister clades to the flatworms (e.g. nemerteans and annelids [2,45,46]. At present, transcriptomic data for an increasing diversity of these taxa is available [47,48], but data on mode of development for many of these species is lacking.

The evolution of larvae within Spiralia and Metazoa is a convoluted and recurring topic. In order to better describe the origins of indirect development, analyses need to include species that span as much variation in mode of development as possible. Curiously it might be the inclusion, and the study, of direct developing species that may reveal interesting findings that aid our interpretation of life history evolution. A vestigial prototroch during embryogenesis in an early branching, direct developing nemertean, for example, suggests that the trochophore larvae was lost in this clade [45].

## CONCLUSION

This study represents the first phylogenomic framework of polyclads and the first ancestral state reconstruction of mode of development in flatworms. Our phylogenomic analysis of polyclads revealed the macroevolutionary distribution of developmental modes across superfamilies, families and genera, and sheds light on the ancestral condition for many clades. It suggests that indirect development has evolved secondarily within, or on the lineage leading to, Polycladida and has likely been gained or lost several times. Our findings support the hypothesis that indirect development (and characters, such as larval ciliary bands) is homologous among prosthiostomid, euryleptid, and pseudocerotid cotyleans, but was unable to resolve homology of indirect development between the polyclad suborders (Cotylea and Acotylea). Homology of indirect development between flatworm orders (Polycladida and Neodermata), and among flatworms and other spiralian lineages, seems unlikely. Alternative scenarios, such as multiple losses of ancestral flatworm larvae, remain possible but are less parsimonious. Increased taxon sampling with robust transcriptomic (or genomic) data will allow for a more detailed understanding of the complex evolution of different larval forms within Polycladida and across flatworms more generally. Polyclad and neodermatan flatworms make excellent systems for understanding how indirect development evolves and larval characters diversify, particularly with regard to the intercalation hypothesis.

## METHODS

### Organismal sampling

One or more specimens of each of the 21 representative species were collected in the intertidal and shallow subtidal, in tide pools or via snorkeling or SCUBA (self-contained underwater breathing apparatus; under AAUS certification) using direct, non-destructive collecting under rocks. *Theama mediterranea* was extracted from sand samples collected near Rovinj, Croatia (see [54]). A visual examination was used for confirmation of identity for fifteen species *Boninia divae* (Hyman, 1955), *Prosthiostomum acroporae, Enchiridium periommatum*, *Idioplana atlantica*, Planoceridae sp., *Phaenocelis* sp. *Phrikoceros mopsus* (Marcus, 1952), *Thysanozoon* sp. or *Acanthozoon* sp., *Pseudobiceros damawan* Newman & Cannon, 1994, *Pseudoceros paralaticlavus* Newman & Cannon, 1994, *Theama mediterranea, Euplana gracilis, Gnesioceros sargassicola, Euprosthiostomum mortensi*, and *Thysanozoon brocchi* (Risso, 1881)). The identification of the other six species was carried out using morphological analysis of histological sections (see methods below). At least one specimen was placed in RNAlater solution (Qiagen, Hilden, Germany) for RNA preservation and frozen at −80 C within one week of collection to prevent RNA degradation. A second specimen of each species, when available, was fixed as a voucher for morphological analysis, first in 4% formalin using the frozen formalin technique [55] and subsequently preserved in 70% ethanol for long-term storage. For histology, specimens in 70% ethanol were graded into 100% ethanol, cleared in Histoclear (National Diagnostics) for 1 h, infiltrated with 1:1 histoclear/paraffin for 24 h and equilibrated in molten paraffin for 24 h (all steps performed at 60 C). Specimens were then embedded in fresh paraffin and left to harden at room temperature for 24 h. Specimens were sectioned in the sagittal plane at 8 μm on a rotary microtome, mounted on glass slides and stained with Masson’s trichrome [56]. Identification to genus level was achieved using the taxonomic monographs of Faubel (1983, 1984) and Prudhoe (1985). Species level ID was achieved by consulting the species descriptions in the literature, and also verified by comparing 28S rDNA sequence data from our transcriptomes to the polyclad 28S rDNA sequences available on NCBI Genbank. Specimens not completely used up by RNA extraction were deposited in the Smithsonian National Museum of Natural History (NMNH) and are available for study under the catalog numbers provided in Table S1.

We generated RNA-Seq data for 21 polyclad species, and downloaded data for six additional polyclad species from the NCBI Sequence Read Archive (SRA). We also obtained data from nine outgroup species from the SRA: four species from Prorhynchida (the sister taxon to Polycladida [11], including *Geocentrophora applanata* (Kennel, 1888), *Prorhynchus alpinus* Steinböck, 1924, *Prorhynchus* sp. I Laumer & Giribet, 2014, and *Xenoprorhynchus* sp. I, three species from the sister taxon of Polycladida + Prorhynchida [11], including *Gnosonesimida* sp. IV Laumer & Giribet, 2014, *Schmidtea mediterranea* (Benazzi et al., 1975), and *Taenia solium* (Linnaeus, 1758), one species from Macrostomorpha (*Microstomum lineare* (Müller OF, 1773)), and one species from Catenulida (*Stenostomum leucops* (Duges, 1828)). Specimen data and Sequence Read Archive (SRA) accession numbers are listed in Table S1.

### RNA extraction and sequencing

A 20–100 mg tissue sample was taken from the anterior of each animal and homogenized using a motorized pestle. In some cases, the specimen was so small the entire animal was used. For *Theama mediterranea*, 20 adults, starved for 1 month, were extracted in a single tube using a protocol detailed in [10]. For all other polyclads, the tissue was homogenized for 1-2 min, then it was flash-frozen in liquid nitrogen for subsequent homogenizing, until tissue mixture was fully uniform. TriZOL Reagent (Life Technologies, Carlsbad, CA, USA), 500 μL, was then added and the mixture was completely homogenized. Once this process was complete, an additional 500 μL of TriZOL Reagent was added to the solution and the mixture was left at room temperature for five min. Following the five min incubation, 100 μL of 1-Bromo-3-chloropropane was added to the solution, which was subsequently mixed thoroughly by vortexing the sample for 10 s. The mixture was then left at room temperature for five min, and then centrifuged at 16,000 g for 20 min at 8 C. The top aqueous phase was then removed and placed in another tube where 500 μL of 100% isopropanol was added, and stored for 1 h at −20 C for RNA precipitation.

After precipitation, the samples were centrifuged at 17,200 g for 10 min at 4 C. The supernatant was then removed and the pellet washed with freshly prepared 75% ethanol. The sample was then centrifuged at 7,500 g for 5 min at 4 C. The supernatant was removed and the pellet air-dried for 1 to 2 min (or until it looked slightly gelatinous and translucent). The total RNA was then re-suspended in 10–30 μL of Ambion Storage Solution (Life Technologies, Carlsbad, CA, USA), and 1 μL of SUPERase•In (Thermo Fisher Scientific, Waltham, Massachusetts, USA) was added to prevent degradation.

Total RNA samples were submitted to the DNA Sequencing Facility at University of Maryland Institute for Bioscience and Biotechnology Research, MD, USA or The Hospital for Sick Children Centre for Applied Genomics in Toronto, ON, Canada, where quality assessment, library preparation, and sequencing were performed. RNA quality assessment was done with a Bioanalyzer 2100 (Agilent Technologies, Santa Clara, CA, USA), and samples with a concentration higher than 20 ng/μL were used for library construction. Library preparation used the Illumina TruSeq RNA Library Preparation Kit v2 (Illumina, San Diego, CA, USA) and 200 bp inserts; 100 bp or 125 bp (*Theama* and *Boninia*), paired-end reads were sequenced with an Illumina HiSeq1000 and HiSeq2000 sequencers (Illumina, San Diego, CA, USA).

### Quality control and assembly of reads

Reads that failed to pass the Illumina “Chastity” quality filter were excluded from our analyses. Reads passing the quality filter were assembled using Trinity (version 2.4.0 for most, but version 2.6.6 for species *Boninia divae* and *Theama mediterranea*; [57]) with default settings, which required assembled transcript fragments to be at least 200 bp in length. Reads were trimmed pre-assembly for the species *Boninia divae* and *Theama mediterranea* using Trimmomatic [58]. Assemblies are available at https://doi.org/10.6076/D1JG60. Assembly quality was assessed using Benchmarking Universal Single-Copy Orthologs (BUSCO) v5.4.2 [59].

### Orthology assignment

Translated transcript fragments were organized into orthologous groups corresponding to a custom platyhelminth-specific core-ortholog set of 9,157 protein models (constructed in the same manner as in [60]) using HaMStR (version 13.2.6; [61]), which in turn used FASTA (version 36.3.6; [62]), GeneWise (version 2.2.0; [63]), and HMMER (version 3.1b2; [64]). In the first step of the HaMStR procedure, substrings of assembled transcript fragments (translated nucleotide sequences) that matched one of the platyhelminth protein models were provisionally assigned to that orthologous group. To reduce the number of highly divergent, potentially paralogous sequences returned by this search, we set the E-value cutoff defining an HMM hit to 1×10^−5 (the HaMStR default is 1.0), and retained only the top-scoring quartile of hits. In the second HaMStR step, the provisional hits from the HMM search were compared to our reference taxon, *Echinococcus granulosus* (Batsch, 1786), and retained only if they survived a reciprocal best BLAST hit test with the reference taxon using an E-value cutoff of 1×10^-5 (the HaMStR default was 10.0). In our implementation, we substituted FASTA for BLAST [65] because FASTA programs readily accepted our custom amino acid substitution matrix (POLY90). Both the Platyhelminthes core-ortholog set and custom substitution matrix are available at https://doi.org/10.6076/D1JG60.

The Platyhelminthes core-ortholog set was generated by first downloading all available platyhelminthes clusters with 50% similarity or higher from UniProt [66] (70,698 clusters). Excluding clusters that contained only one sequence left 20,874 clusters. We calculated the sequence similarity of each cluster and as a heuristic, decided to remove clusters whose percent identity was less than 70%, which left 20,549 clusters. We then assessed the number of times each taxon was represented within those clusters. *Echinococcus granulosus* was identified as the most closely related, most abundant taxon (9,157 associated clusters with 70% similarity or higher), and was therefore selected as the reference taxon for the custom HaMStR database. We constructed the platyhelminthes HaMStR database by following the steps given in the HaMStR README file, which included generating profile hidden Markov models for each cluster using HMMER. Our platyhelminthes HaMStR database contained 9,157 orthologous groups. All protein sequences for *Echinococcus granulosus* (UniProt/NCBI taxon ID 6210) were downloaded from UniProt and used to generate the BLAST database for HaMStR.

Construction of the custom substitution matrix (PLATY90) followed the procedure outlined in Lemaitre et al. [67], which used only greater than 90%-similarity platyhelminthes clusters downloaded from UniProt with singleton clusters removed. Use of a taxonomically-focused amino acid substitution matrix follows similar procedures used in arthropods [68] and gastropods [60,69] that seek to improve the amino acid alignments performed in the process of a phylogenomic workflow. In this protocol, a block is defined as a conserved, gap-free region of the alignment. Our blocks output file contained 205,562 blocks.

### Construction of data matrix and paralogy filtering

Protein sequences in each orthologous group were aligned using MAFFT (version 7.187; [70]). We used the --auto and --add fragments options of MAFFT to align transcript fragments to the *Echinococcus granulosus* reference sequence, which was considered the existing alignment. We converted the protein alignments to corresponding nucleotide alignments using a custom Perl script. A maximum likelihood tree was inferred using RAxML-NG (RAxML Next Generation version 0.6.0; [71]) for each orthologous group where at least 75% of the taxa were present (4668 orthologous groups), and was given as input to PhyloTreePruner (version 1.0; [72]). Orthologous groups that showed evidence of out-paralogs for any taxa (2530 orthologous groups out of 4426) were pruned according to the default PhyloTreePruner protocol, which removes all additional sequences outside of a maximally inclusive sub-tree. For orthologous groups containing in-paralogs, multiple sequences were combined into a single consensus sequence for each taxon, and orthologous groups for which fewer than 75% of taxa remained were discarded. This process left 4469 orthologous groups eligible for inclusion in our data matrices. Individual orthologous group alignments were concatenated, and codons not represented by sequence data in at least four taxa were then removed.

### Phylogenetic analyses

For phylogenetic analysis, the final nucleotide data matrix from transcriptome data was partitioned by codon position by assigning different model parameters and rates to the three codon positions. We conducted the phylogenetic analysis using RAxML-NG (version 0.6.0; [71]). We used the default settings in RAxML-NG, and partitioned our data set by codon position. Each partition was assigned a general time reversible substitution model (GTR; [73]) with a rate heterogeneity model with a proportion of invariant sites estimated (+I) and the remainder with a gamma distribution (+G; [74]]), along with stepwise-addition starting trees. For our analysis, 500 bootstrap replicates were generated and a best tree search was performed with 20 search replicates. To assess whether heterotachy may be impacting our inferences, we also ran a maximum likelihood analysis with IQ-TREE v2.2.0 [75] under the GHOST model of evolution [76] with ultrafast bootstrap approximation [77]. Data matrices and phylogenetic analysis outputs are available at https://doi.org/10.6076/D1JG60.

### Ancestral state reconstruction

Ancestral states were reconstructed for development type (indirect or direct) for the complete polyclad phylogeny plus outgroups (Table S3). We assessed fit for two models using the corrected AIC (AICc), where: (i) all transition rates were equal (ER; same as the symmetrical model in this case); (ii) forward and reverse transitions were different between states (all rates different, ARD). The ER model (AICc = 24.77317) was a slightly better fit to the data than the ARD model (AICc = 25.86218 for development type). In order to more confidently infer ancestral states across the phylum Platyhelminthes, we also reconstructed ancestral development type for the whole phylum using a previously published phylogeny [11]. The goal of this reconstruction was to reduce bias caused by increased sampling of polyclad species compared to other groups in our phylogeny. Tree manipulation was conducted using the APE package [78], and the final ancestral state reconstruction analysis and model testing was completed using the rayDISC function in the corHMM package [79]. The package corHMM fits a hidden rates model that treats rate classes as hidden states in a Markov process, employing a maximum likelihood approach. When a state is missing for a particular species, RayDISC assigns equal likelihoods to both states (indirect or direct development). In this analysis, the marginal ancestral states are returned, which are given as the proportion of the total likelihood calculated for each state at each node. To test for significance of ancestral state reconstruction, we used proportional likelihood significance tests under the rule of thumb that a log likelihood difference of 2 or greater represents a significant difference. R-scripts are available at https://github.com/goodgodric28/polycladida_phylogenomics.

## ETHICS

Specimen collection was conducted under the auspices of the Florida Fish and Wildlife Conservation Commission (Special Activity Licence no. SAL-14-1565-SR), the Délégation Régionale à la Recherché et à la Technologie de la Polynesie Francaise (no number for permit), and the Curaçaoan Government (permits provided to The Caribbean Research and Management of Biodiversity foundation; Government reference: 2012/ 48584). Sampling and using *Theama mediterranea* in this study was permitted according to the Croatian Nature Protection Act published in the Official Gazette No. 80/2013.

## DATA ACCESSIBILITY

Transcriptomes can be accessed at the SRA at NCBI: SRA accession numbers SRR15530025– SRR15530044 (Table S1). Critical files from our phylogenomics pipeline are available on DataDryad (https://doi.org/10.6076/D1JG60). Scripts and files from our ancestral state reconstruction analyses are available on Github (https://github.com/goodgodric28/polycladida_phylogenomics).

## AUTHOR CONTRIBUTIONS

JAG, KAR AGC and MPC conceived of and designed the study; JAG, BE and KAR collected samples and field data; JAG and BE carried out the molecular lab work; JAG and BE performed the bioinformatics analyses; all authors participated in data analysis, helped draft the manuscript, and gave final approval for publication.

## COMPETING INTERESTS

We have no competing interests.

## FUNDING

This research was supported by the Global Genome Initiative under Grant No. GGI-Rolling-2016-035, NSF Partnerships for International Research and Education program Award 1243541. JAG was supported by the Smithsonian Institution (Peter Buck Pre-doctoral Fellowship) and the NSF (PRFB-1711201). BE was supported by a grant for young scientists by the University of Innsbruck.

## ACKNOWLEDGEMENTS

We thank Ariane Dimitris, Joie Cannon, Kevin Kocot, Isabel Dittmann, Raimund Schnegg, and Rebecca Varney for their collecting assistance. We are also grateful to The Caribbean Research and Management of Biodiversity foundation (CARMABI) in Curaçao, the Smithsonian Marine Station at Fort Pierce, Florida USA, and the Richard B. Gump South Pacific Research Station in French Polynesia for use of their facilities and assistance in acquiring the proper permits. Finally, we thank the Laboratories of Analytical Biology of the National Museum of Natural History for the use of their laboratory facilities.

## SUPPLEMENTARY TABLES

**Table S1.** Samples included in our polyclad phylogenetic reconstruction (Figure 1). Table includes SRA accession numbers, collecting locality, and method of identification.

**Table S2.** Transcriptome assembly and orthology assignment statistics for samples included in our polyclad phylogenetic reconstruction (Figure 1).

**Table S3.** Mode of development for species and genera in this study recorded from the literature and personal observation.

## SUPPLEMENTARY FIGURES

**Figure S1.** Histological sections showing the male and female reproductive structures used to identify some of the polyclad taxa in this study. A) Posterior end of *Eucestoplana* sp. showing female and male openings (anterior right), B) *Cycloplorus gabriellae*, i) female gonopore and male seminal vesicle (asterisk) and penis (arrow), ii) prostatic vesicle (PV)(anterior left). C) *Pericelis cata* showing sucker (arrowhead) behind female aperture (arrow)(anterior left). D) *Notocomplana* sp. (anterior left) showing seminal vesicle (asterisk), interpolated prostatic vesicle (PV) with tubular chamber, and female opening, E) *Notocomplana lapunda* (posterior left) showing unarmed penis (arrow) and Lang’s vesicle (asterix). F) *Armatoplana leptalea* (posterior left) showing penis armed with long, sharp stylet housed in a tubular atrium (arrow) and seminal vesicle (asterisk). G) *Phaenocelis* sp. (posterior left) showing prostatic vesicle (PV) (i), rod-like penis (arrow) (ii), and Lang’s vesicle (iii)(asterisk). Scale = 500um.

## Notes

https://doi.org/10.6076/D1JG60

https://github.com/goodgodric28/polycladida_phylogenomics

https://www.ncbi.nlm.nih.gov/bioproject/PRJNA544137

### Competing Interest Statement

The authors have declared no competing interest.

### Summary of Updates

We have revised the title and text of the manuscript to highlight that our tree does not address whether polyclad larvae are homologous to the trochophore larvae of other lineages. We have provided more information in the methods, results, and discussion to explain our choice to include a secondary, order level, reconstruction. The primary reasoning is that although species-level reconstructions are generally preferred, our phylogenomic analysis of polyclads is heavily biased towards polyclad species and therefore was not sufficient to reconstruct earlier ancestral nodes. The addition of a previously published Platyhelminthes tree allowed us to maintain a more balanced and thorough sampling across the different Platyhelminthes lineages in order to determine that polyclad larvae are likely secondarily derived. BUSCO scores have been added to Table S2 with a summary in the results section. We did not previously model heterotachy in our analysis. However, a model of evolution was recently published (GHOST; https://pubmed.ncbi.nlm.nih.gov/31364711/) that accounts for heterotachy in ML phylogenetic analyses (implemented in IQ-Tree). We have added this analysis to our manuscript. The resulting tree is identical with regard to relationships among taxa and is nearly identical in terms of branch lengths, so we have left the original analyses untouched.

