## Supplementary figures and images for "A phylogenomic approach to resolving interrelationships of polyclad flatworms, with implications for life history evolution"

### Figure S1

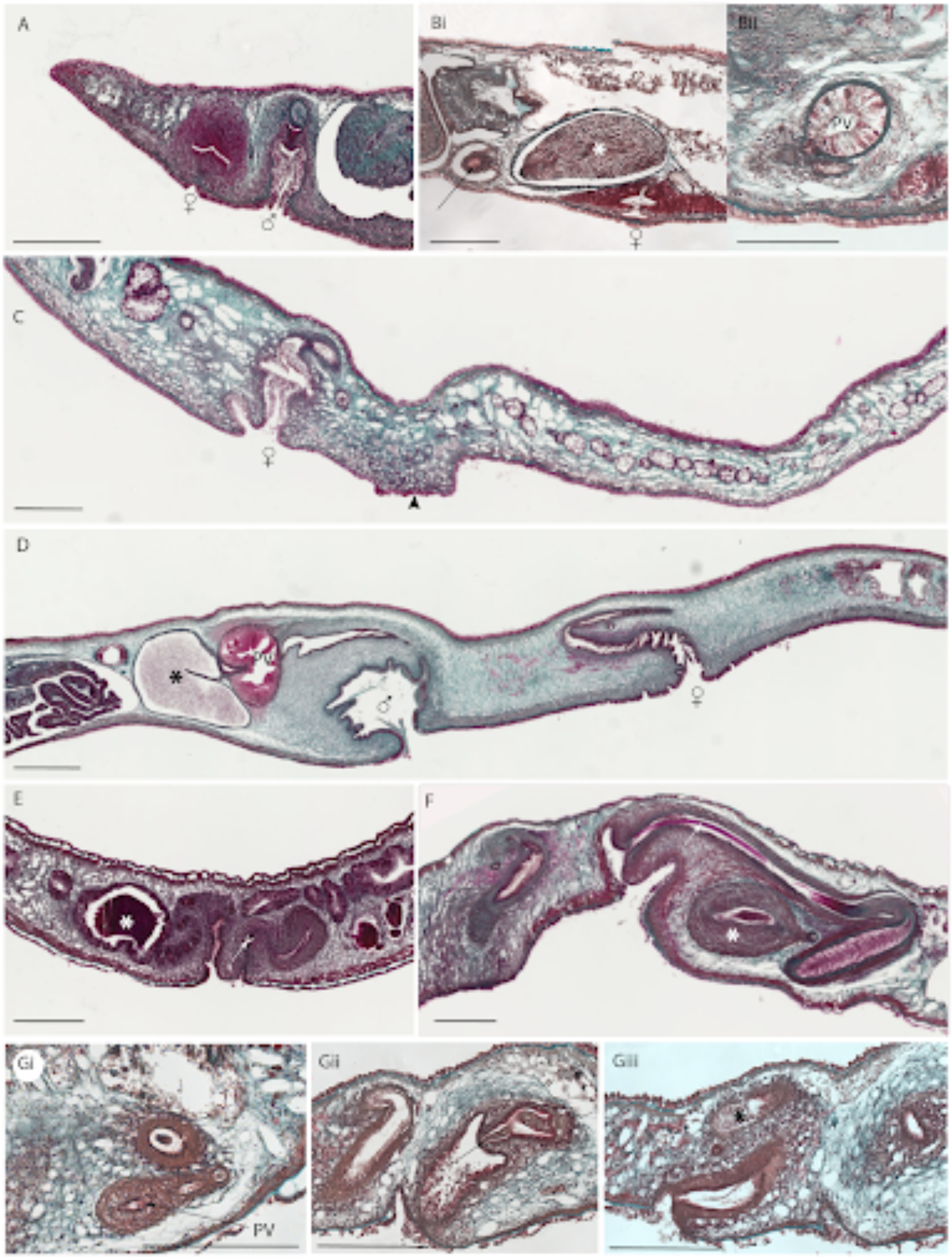
